# *µPIX*: Leveraging Generative AI for Enhanced, Personalized and Sustainable Microscopy

**DOI:** 10.1101/2024.10.25.620201

**Authors:** Gabriel Bon, Daniel Sapède, Cédric Matthews, Fabrice Daian

## Abstract

Fluorescence microscopy is a critical tool in bio-cellular research, enabling the visualization of biological tissues and cellular structures. However, the inevitable aging of microscopes can degrade their performance posing challenges for long-term scientific investigations. In this study, we introduce *µPIX*, a personalized deep learning workflow based on a Generative Adversarial Network (GAN) utilizing a Pix2Pix architecture. The network is trained in a supervised manner to denoise images, optimize pre-processing for binary segmentation, and compensate for equipment aging. Our results, evaluated using standard image quality and binary segmentation metrics, demonstrate that *µPIX* outperforms popular deep learning architectures based on convolutional auto-encoder networks for similar tasks. Additionally, our generative model effectively rejuvenates older detectors to perform on par with newer ones, not only by improving image quality but also by preserving resolution in depth and maintaining a near-linear response between original and generated images in terms of pixel intensity (crucial for quantitative imaging). These findings suggest that generative deep learning approaches can significantly contribute to more sustainable, cost-effective microscopy, fostering continued innovation and discovery in biological research.

## INTRODUCTION

Microscopy is an indispensable tool in biological research, enabling the visualization of structures at cellular and molecular levels. However, the performance of microscopes degrades over time leading to diminished signal-to-noise ratio (SNR), reduced resolution and other artifacts that impair image quality. These issues necessitate frequent maintenance that is generally not readily available, hardware upgrade and eventually, replacement of expensive microscopy equipments. In an era where sustainability and cost-effectiveness are paramount, extending the functional lifespan of existing microscopy systems is both economically and environmentally beneficial. The aging of microscopes manifests in several ways, including decreased light throughput, increased background noise, and deteriorating optical alignment. These factors collectively reduce the SNR, making it challenging to discern fine details and thus make accurate quantitative measurements on images of biological samples. Historically, image restoration in fluorescence microscopy relied on techniques like deconvolution and filtering. Classical methods like Wiener filtering and Richardson-Lucy deconvolution have been widely used to reduce noise and correct blur (1; 2; 3; 4). However these approaches often require precise knowledge of the point spread function (PSF) and can be computationally intensive, limiting their applicability to real-time imaging scenarios or are expensive when supplied by manufacturers (5). This solution allows action to be taken in the time interval between the progressive loss of signal, due in particular to the loss of detector efficiency, and the moment a fault occurs resulting in an inoperative system.

In recent years, Deep Learning has revolutionized the field of image restoration. Convolutional neural networks (CNNs), GAN and more recently Diffusion models have demonstrated their superior performance in image denoising, super-resolution, and artifact removal compared to traditional methods (6). For example, the Content-Aware Image Restoration (CARE)(7) networks use a classical U-Net architecture in the context of a convolutional auto-encoder to restore images by learning from pairs of low-quality and high-quality images, real or synthetic. This approach has proven effective in reducing noise and enhancing resolution without requiring PSF information. Noise2Void (8) and Noise2Noise (9) are notable techniques that further simplify the training process by eliminating the need for training paired data in an unsupervised manner. CellPose (10; 11), a generalist deep learning algorithm offers a robust image restoration capabilities alongside its primary function of cell segmentation. In its last version 3, it includes specialized models for denoising, debluring and upsampling, tailored for both cytoplasmic and nuclear channels (12). This flexibility allows it to handle a wide range of image degradation issues and offers an accurate segmentation of cells. The W2S framework (13; 14) combines wavelet transformations together with CNNs to enhance both the resolution and the contrast of fluorescent microscopy images. This method leverages multi-scale information to improve image quality. Similarly, the Deep-Z framework (15) enables virtual refocusing in 3D fluorescence microscopy, significantly extending the depth of field and correcting optical aberrations. High-throughput imaging of 3D samples, such as tumor spheroids and organoids has also benefited from Deep Learning. Techniques that combine axial z-sweep image acquisition with CNN based restoration allow for faster imaging with reduced photo-toxicity, crucial for live imaging. These methods can generate high-quality 2D projections from low-quality z-sweep images enabling real-time analysis with minimal exposure times.

Despite these advancements, challenges remain in generalizing deep learning models across different imaging conditions and microscopy setups. Our proposed workflow leverages these advancements to address both immediate and long-term challenges in microscopy image restoration. In contrast to current state-of-the-art model based on UNET architectures (16), our *µPIX* workflow leverages the use of a conditional generative adversarial network (cGAN) to tackle efficiently the classical denoising and segmentation problematics. Moreover, our approach allows us to tailor a highly specialized model by implementing a precision-focused strategy ensuring that our enhancements are optimally aligned with the specific deficiency and operational context of the equipment leading to superior image restoration and extended utility of the microscope.

## RESULTS

### *µPIX* is built on a generative Pix2Pix architecture

Image denoising is known to be one of the major problems in the field of image analysis and deep Learning based solutions have proven their superior capabilities in this task in comparison to traditional denoising algorithms. As of today, the architecture based on convolutional autoencoder are considered as the state of the art to tackle this problem. UNET and similar convolutional autoencoders operate primarily through pixel-wise predictions, optimizing pixel-level accuracy using reconstruction loss functions. While this approach ensures a good and fast overall image reconstruction, it can struggle with preserving fine details and textures, especially in denoising tasks. In this context, we first based our approach on the use of a classical UNET network working in combination with a classical pixel-wise loss (mean squared error - MSE) and a perceptual loss network (VGG16) to make our network aware of high level perceptual and semantic differences between original and predicted images. Unfortunately, even if we improved slightly in image reconstruction quality, in comparison to classical UNET networks used for such tasks, the results were not satisfying. To address these limitations, we chose to base our workflow on Pix2Pix generative network ***(Figure 1)***. Unlike UNET, *µPIX* leverages the cGAN architecture to produce high-quality and realistic denoised images. This choice allows us to train a model that is not only capable of tackling classical image challenges like denoising but also able to address the specific defects of particular hardware. By leveraging the flexibility of the Pix2Pix architecture, *µPIX* can adapt to the unique characteristics and imperfections of specific imaging equipment, effectively building a specialized prosthesis for each device. This adaptability ensures that our model can provide optimized solutions tailored to the nuances of different hardware, enhancing overall performance and image quality in a way that traditional convolutional autoencoders cannot achieve. Our network introduce an adversarial loss that encourages the network to generate images that are not only accurate but also perceptually convincing and realistic. This adversarial training helps preserve fine details and textures that are often lost in pixel-wise approaches. The network is compounded of two subnetworks working together: a generator and a discriminator. The generator network is trained to learn how to transform an input image to resemble a reference image provided during the supervised training. Classically this subnetwork architecture is based on a classical UNET network. The encoding part of the generator has been chosen carefully as we wanted to keep a good compromise between training/inference speed and performance. We decided to choose a lightweight and performant EfficientNet-b0 (17) as the backbone for this subnetwork. To ensure that the training phase will be efficient and will not collapse quickly, we decided to not use a pre-trained version of this backbone to avoid at start a too big gap between generator and discriminator capabilities. The discriminator subnetwork is based on a PatchGAN network derived from a classical convolutional neural network which has proven its superior discrimination capabilities in such architectures (18). The main objective of the discriminator is to assess whether the given images as input are generated or real images. Through adversarial training (19), in a supervised manner, the generator and discriminator iteratively improve their accuracy, leading to the generator becoming increasingly accurate at producing images that closely resemble the reference images. At the end of theses steps, the inference phase will only use the generator subnetwork for image generation.

**Figure 1.**
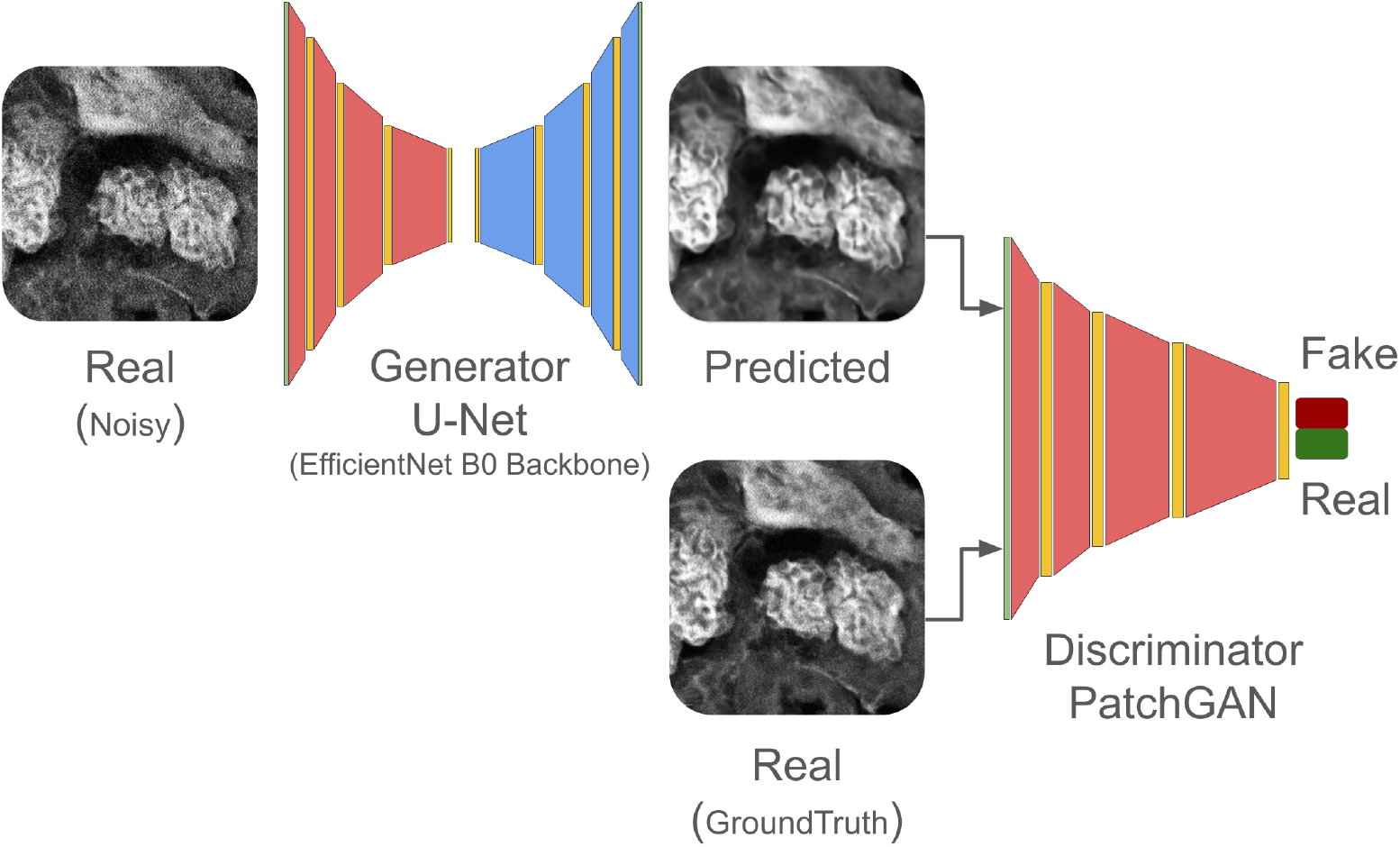
*µPIX* architecture is based on a Pix2Pix generative network. *µPIX* consists of two subnetworks: a generator, based on a UNet architecture with an EfficientNet-b0 backbone, and a discriminator (PatchGAN). During supervised training, a noisy image is input to the generator, which produces a predicted image. This output is compared to the real clean image using a pixel-wise loss function (MSE). Pairs of real and generated images are then passed to the discriminator, which classifies them as real or fake using a binary cross-entropy loss (BCE). Both subnetworks are progressively refined through adversarial loss during training. In the inference phase, only the trained generator is used to produce clean images.

### *µPIX* outperforms state-of-the-art denoisers on CSBDeep Denoising Benchmark Dataset

Image denoising and restoration are central challenges in the analysis of microscopy data. To assess the accuracy of our workflow, we chose to use the CSBDeep Denoising Dataset as a benchmark dataset to evaluate our approach (7). This dataset, derived from the Broad Bioimage Benchmark Collection (20), comprises pairs of clean reference images containing cell nuclei and their corresponding artificially noised counterparts from the human U20S cell line. The noising involves synthetically adding significant read-out and shot noise, along with additional 2×2 pixel binning to mimic acquisitions at very low light levels. To evaluate our workflow comprehensively, we used a set of denoising and image restoration quality metrics. We employed the mean-squared error (MSE) as a measure of signal-to-noise ratio by comparing the reference clean images to the images generated by *µPIX* from the noisy image. The Structural Similarity Index (SSIM) was used to measure the preservation of overall object shapes. We compared our approach against two popular denoising software based on UNET architecture: CARE(7) and the “Denoise Nuclei” model from CellPose3 (12). To avoid any generalization and comparison bias, we train CARE from scratch, using the same dataset, and we used the pre-trained version of the Denoise Nuclei from CellPose3, as currently, there is no available method to train this denoiser from scratch using one’s own data. Finally, we evaluated theses architectures and our approach concurrently on the same test dataset consisting of various nuclei images extracted from the original benchmark dataset and we averaged the metrics. As shown in ***Table 1*** and ***Table 2***, our approach clearly outperforms CARE and CellPose3 in terms of both signal-to-noise ratio and structural preservation. For CARE, we improved MSE by 43% and SSIM by almost 11% (MSE: 96.32, SSIM: 0.9002). This superiority can be attributed to several factors: the adversarial process helps the generator produce more realistic and high-quality images, the conditioning provides more context-aware denoising capabilities, and the adversarial loss, compared to pixel-wise loss, encourages the generation of sharper and more structurally accurate images.

**Table 1.**
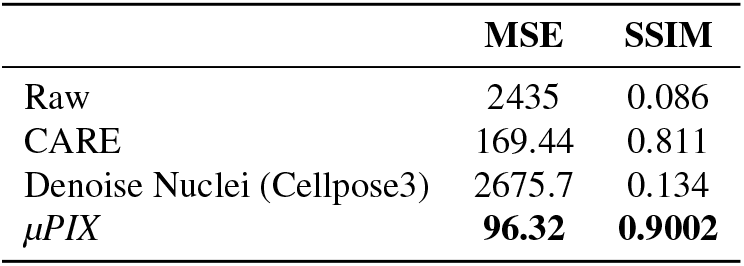
MSE and SSIM metrics for evaluating the quality of denoised images. Three image denoising architectures were used: CARE, Denoise Nuclei, and *µPIX*. For each of these, the MSE and SSIM between the noisy and denoised images from the CSBDeep Denoising Benchmark Dataset test set were evaluated. For reference, the “Raw” row displays the values obtained between the noisy image and the clean (ground truth) image from the test set.

**Table 2.**
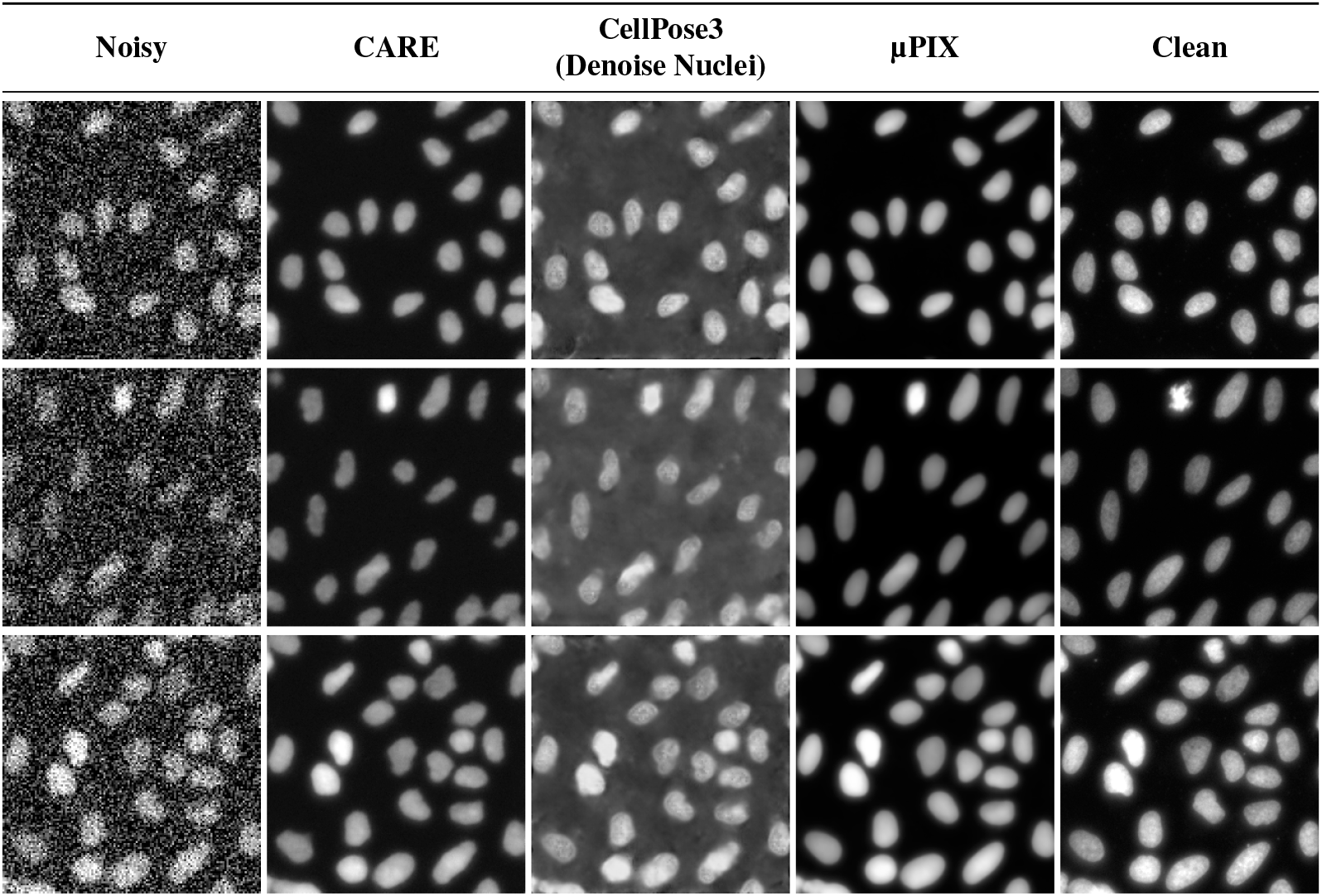
*µPIX* denoising results on CSBDeep Benchmark Dataset. Noisy input images from the CSBDeep Dataset compared to ground truth (clean) images, CARE, CellPose3 Denoise Nuclei, and *µPIX*.

### *µPIX* improves binary segmentation quality when used as an image denoiser in preprocessing steps

Object segmentation is another fundamental task in image analysis and a good preprocessing of images can greatly enhance the further binary segmentation performance. We then wanted to assess how performs *µPIX* as the main image denoiser in a segmentation workflow ***(Figure 2)***. We chose to evaluate *µPIX* against both CARE and Cellpose3 “Denoise Nuclei” as preprocessing steps to enhance image segmentation. As we did not have segmentation ground truth included in the CSBDeep Denoising Dataset and given its robustness and superior performance in segmenting nuclei, we relied on Stardist (21) as image segmenter, using its pre-trained model “2D versatile fluo” to infers binary segmentation from clean images which served as the reference for perfect segmentation. Using the same test dataset previously described, we assessed the impact of different preprocessing methods on binary segmentation performance using classical segmentation metrics: Intersection Over Union (IoU), Precision, Recall, and F1-score. IoU measures the overlap between predicted and true segmentations, offering a direct measure of segmentation accuracy. In the context of binary segmentation, Precision and Recall are crucial for understanding the balance between over-segmentation and under-segmentation. Precision indicates the proportion of true positive results among all positive predictions, thereby reflecting the extent of over-segmentation due to false positives. Recall indicates the proportion of true positives among all actual positives, highlighting the degree of under-segmentation caused by false negatives. The F1-score, as the harmonic mean of Precision and Recall, provides a single metric that balances both aspects, ensuring a robust evaluation of binary segmentation performance. As shown in ***Table 3*** and ***Table 4***, using *µPIX* as a preprocessing step for segmentation outperforms the other methods across all metrics, yielding the best results in terms of IoU (0.861), Recall (0.903), and F1-score (0.9253) on the test dataset. Although CARE achieved the highest Precision score (0.9537), *µPIX* maintained a strong balance with its high Recall. We then assessed whether our approach could be generalized to other state-of-the-art segmentation tools. We used the Cellpose3 segmenter as a reference and employed CARE, Cellpose3 “Denoise Nuclei”, and *µPIX* as preprocessing steps to benchmark segmentation quality using the same metrics and test dataset. As shown in ***Table 3*** and ***Table 4***, *µPIX* yielded again the best overall results across the defined segmentation metrics (IoU: 0.861, Recall: 0.903, and F1-score: 0.9253). Additionally, switching from Cellpose3 “Denoise Nuclei” to *µPIX* within the Cellpose3 workflow improved the F1-score by nearly 3% meaning that the actual CellPose3 workflow could be improved by using a trained *µPIX* as denoiser. Consequently, *µPIX* is not only a strong candidate for pure image denoising but also enhances performance when used in conjunction with state-of-the-art models in the context of image binary segmentation.

**Table 3.**
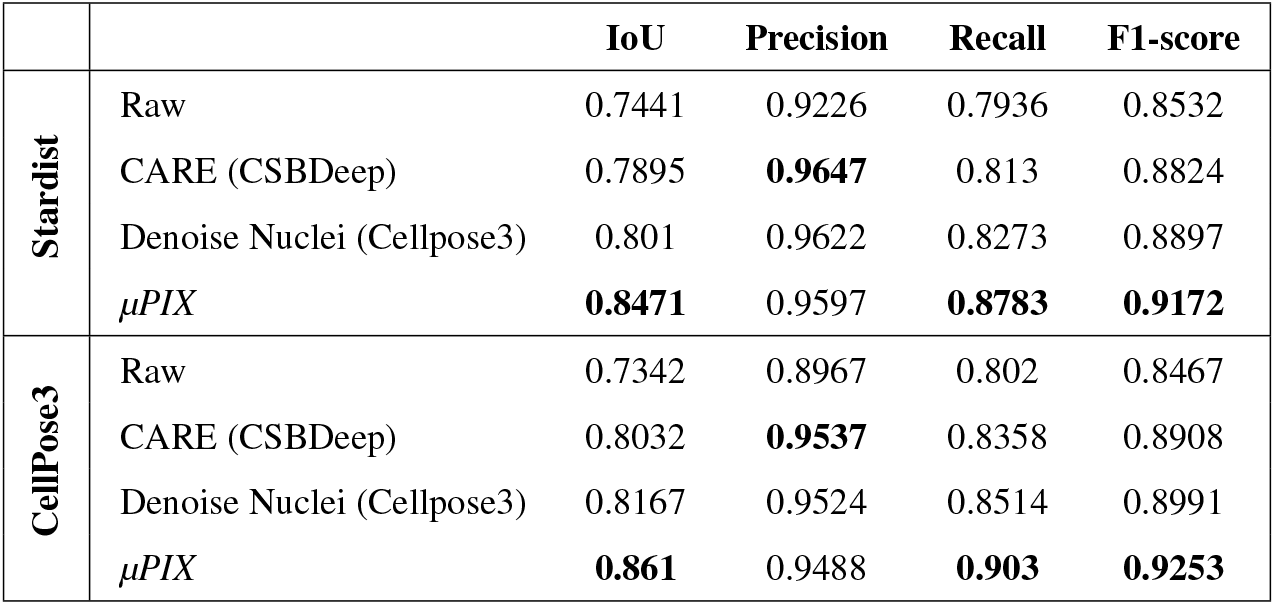
Segmentation metrics using Stardist and CellPose3 as Segmenter. Noisy images from the CSBDeep Benchmark Dataset were denoised using CARE, Denoise Nuclei, and *µPIX*. The denoised images were then segmented using either StarDist or CellPose3. Segmentation metrics (IoU, Precision, Recall, F1-score) were calculated by comparing the binary segmentations of clean and denoised images for both StarDist and CellPose3.

**Table 4.**
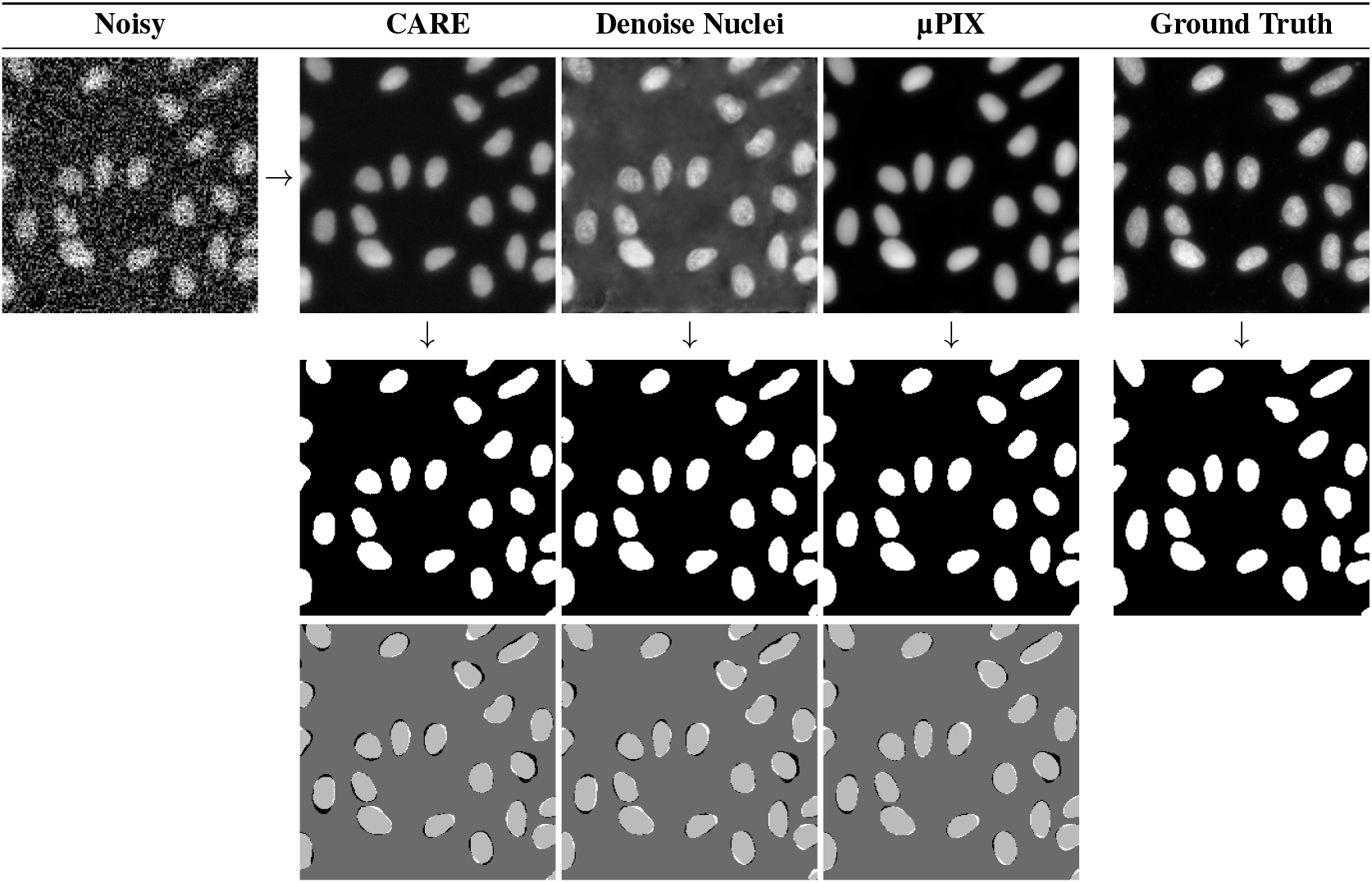
Binary segmentation results on the CSBDeep Benchmark Dataset. **(Top)** Three noisy input examples from the CSBDeep Dataset, along with their denoised counterparts generated by CARE, Denoise Nuclei, and *µPIX*, followed by the ground truth. **(Middle)** Binary segmentation results alongside the ground truth mask. **(Bottom)** Visual representation of segmentation differences: in white false positive, in black false negative and in light gray true positive and dark gray true negative

**Figure 2.**
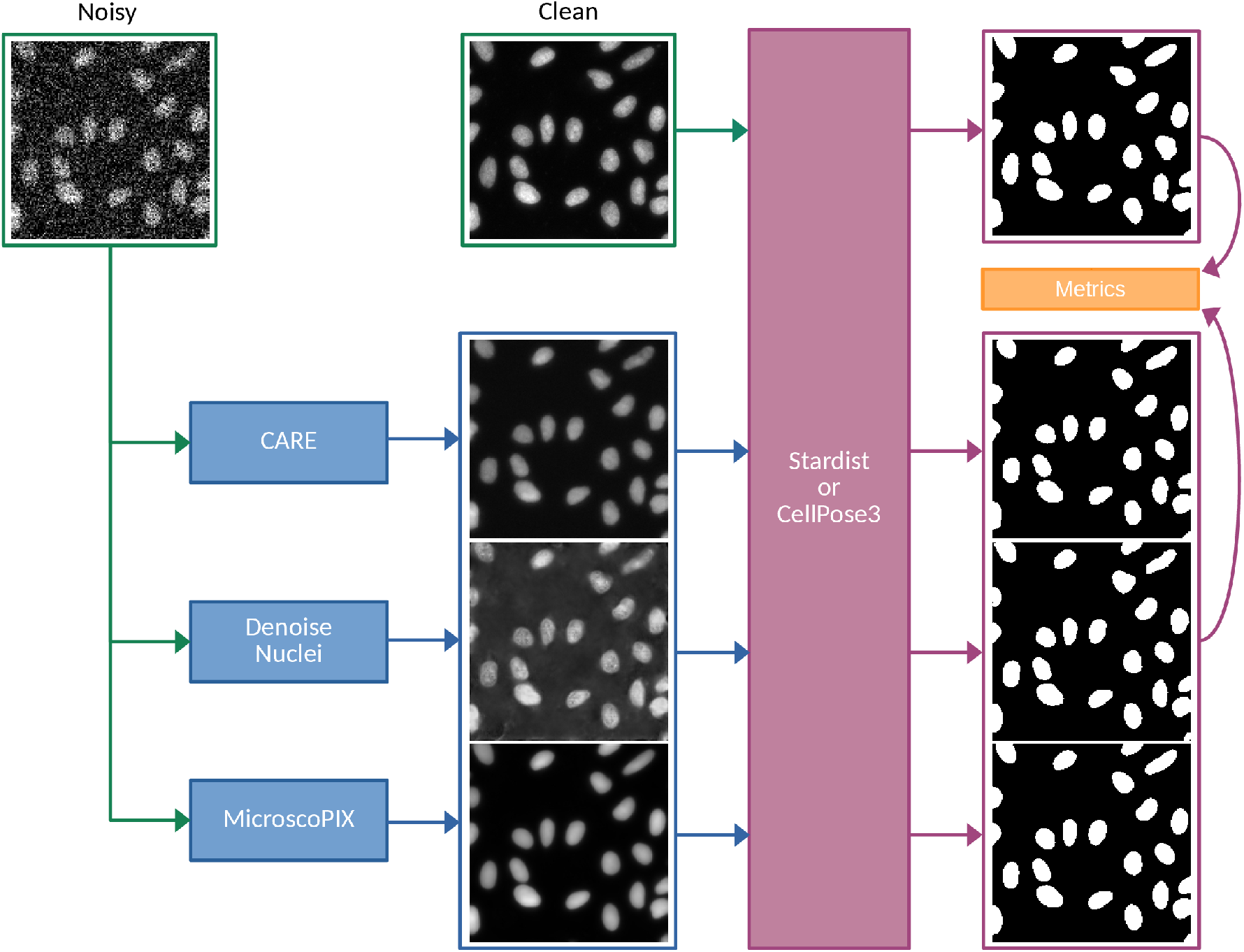
Workflow describing the evaluation process for denoising performance in binary segmentation. To assess the quality of binary segmentation after image denoising by various softwares, pairs of noisy and clean images are used. The noisy images are processed through each of the denoising algorithms: CARE, Denoise Nuclei, and *µPIX* (in blue). The denoised images are then fed into either the Stardist or CellPose3 segmentation models (purple). In parallel, the clean image is also processed by these segmentation models (purple). The resulting binary mask from the clean image serves as the ground truth and is compared with the binary masks generated from the denoised images (orange). The IoU, Precision, Recall, and F1-score metrics are then calculated for performance evaluation.

### *µPIX* enables effective rejuvenation of microscope detectors

A major challenge faced by all microscopy platforms is the aging of equipment, which inevitably introduces acquisition artifacts, making it increasingly complex to analyze acquired data. To demonstrate *µPIX* capabilities toward the challenge of hardware senescence, we chose to simulate one specific case: *detector rejuvenation*. To this end, as we based our architecture on a supervised approach, we chose to simulate detector degradation by building a dedicated dataset consisting of pairs of images acquired simultaneously using an older Multi-Alkali detector (22) and its newer and performant counterpart using a GaAsP detector (23). We acquired this tailored dataset using a biphoton system able to generate two images at the same position within the sample as illustrated in ***Figure 3***. This microscopy setup freed us from the tedious steps of image re-alignment during the acquisition and the dataset preprocessing steps. Furthermore, it allowed us to maintain a supervised learning context mandatory for our Pix2Pix architecture. As we wanted to be as close as real use cases we decided to use biological samples consisting of Gastruloids (24). We acquired two complete stacks, one serving as a training/validation set and the other one as a test set (see Methods). We trained *µPIX* from scratch on this training dataset taking as input the image acquired with the Multi-Alkali detector and considering the corresponding image acquired with GaAsP detector as ground truth. To conclude whether the detector rejuvenation was effective, we decided to check for three different features on image preservation: image quality, intensity preservation along the Z-axis, linear preservation of pixel intensity level between the original and rejuvenated detector. We used MSE and SSIM metrics between images acquired with GaAsP detector and predicted images to assess the quality of the detector rejuvenation. We chose to compare our approach only to CARE because, as of today, there is no published way to train or fine tune a Cellpose3 “denoise nuclei” model on its own dataset.

**Figure 3.**
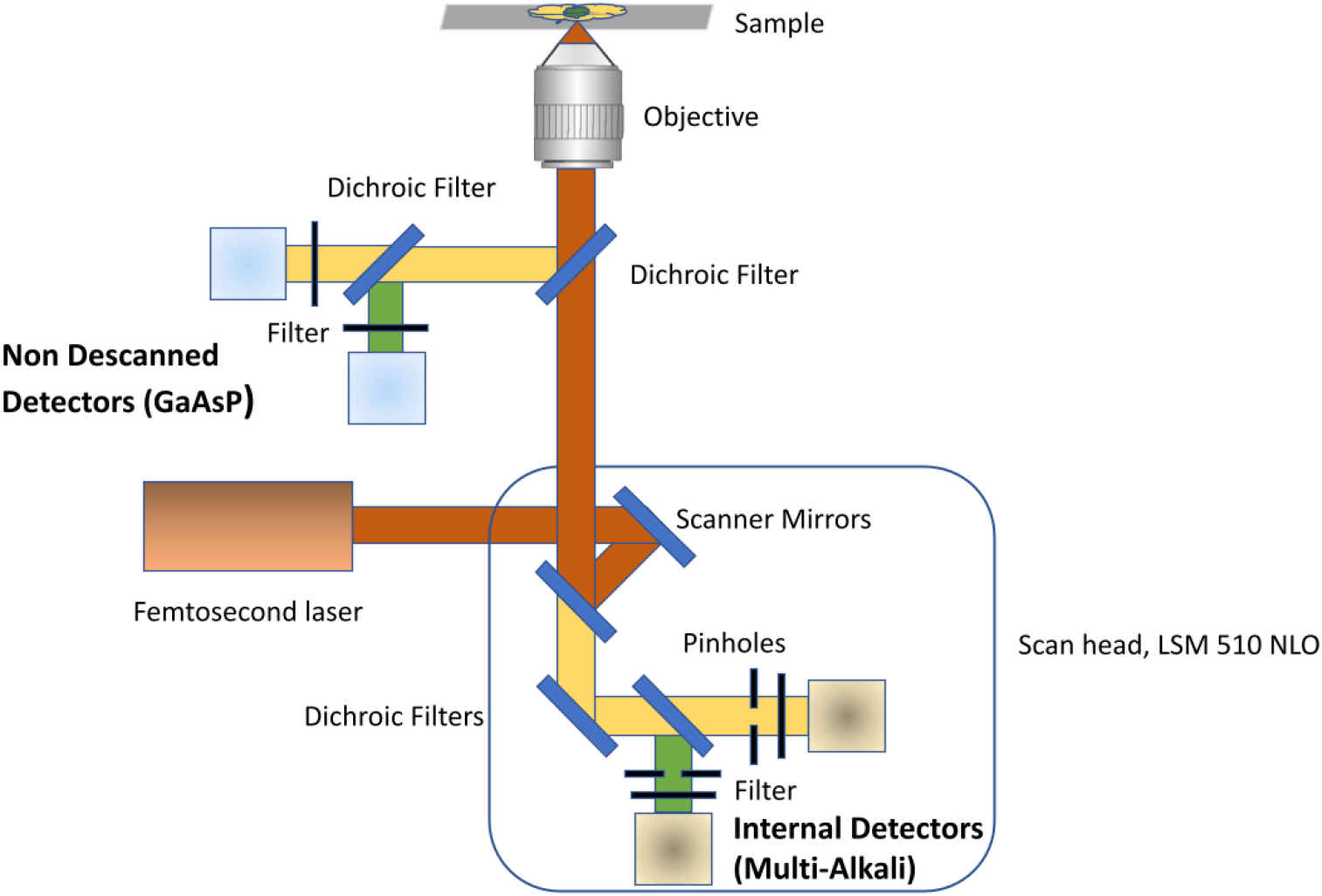
Schematic of a two-photon microscope optical path. Shown here are the two detection channels for the fluorescence signal generated by two-photon absorption. The Non Descanned channel, positioned closest to the objective, is the most sensitive. The GaAsP detectors are new and provide a reference signal optimized for learning. The detection path through the scan head is generally less efficient, even if the pinholes are completely open to collect the scattered emission photons. Multi-Alkali detectors are functional but obsolete, over 15 years old, and will be used to detect ground truth. The signals detected on both types of detectors are almost spatially aligned.

As shown in ***Table 4***, our approach greatly improve the SNR (MSE: 279) and the quality of structure restoration (SSIM: 0.67). Moreover, we can see that *µPIX* successfully handled challenging restoration tasks, such as delineating detected objects, managing complex structures, addressing intense contrast variations (both low and high), and reconstructing objects even in cases where limited information was available ***(Table 5)***,***(Supplementary Table 1)***. These results demonstrate that our approach is effective and reliable for enhancing image quality by compensating the Multi-Alkali detector aging in generating images resembling to ones acquired with a GaAsP detector.

**Table 5.**
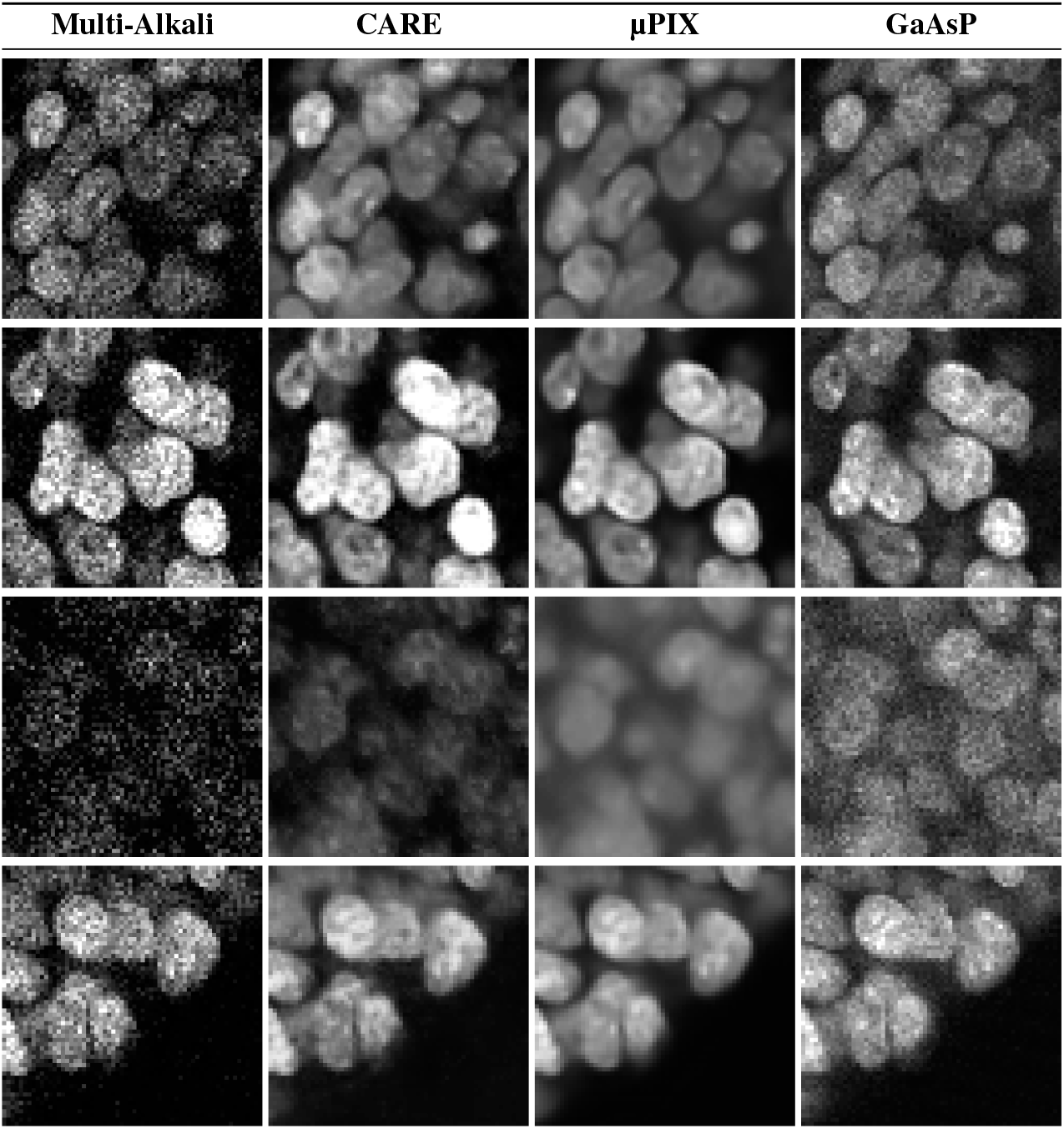
Denoising results on Microscope Rejuvenation Test Dataset. Images acquired using the Multi-Alkali detector from different regions with varying pixel intensities were processed by CARE and *µPIX*, and then compared to the original images obtained with the GaAsP detector.

Moreover, as it is well known that the signal quality diminishes along the Z-axis during a confocal acquisition due to light scattering, light absorption, refractive index mismatch, photo-bleaching and detector sensitivity, we wanted to assess how *µPIX* compensates for these artifacts. To do so, we started by measuring these effects by comparing the MSE between an older Multi-Alkali and a newer GaAsP detector for every slice along the Z-axis of the test stack. As shown in ***Figure 4A***, there is a quasi-exponential difference in terms of MSE as we go deeper into the stack along the Z-axis. This is explainable by the fact that Multi-Alkali detectors are more prone to the signal intensity attenuation effect compared to newer GaAsP detectors. We then compared our predicted images to GaAsP and we see that the difference is far more prone than before and quasi-linear in terms of MSE and SSIM ***(Figure 4B)*** as we go down into the stack along the Z-axis. This means that even if our approach is not able to abolish completely these artifacts, our results in terms of signal intensity are really close and resemble to those we would have obtained using GaAsP detectors.

**Figure 4.**
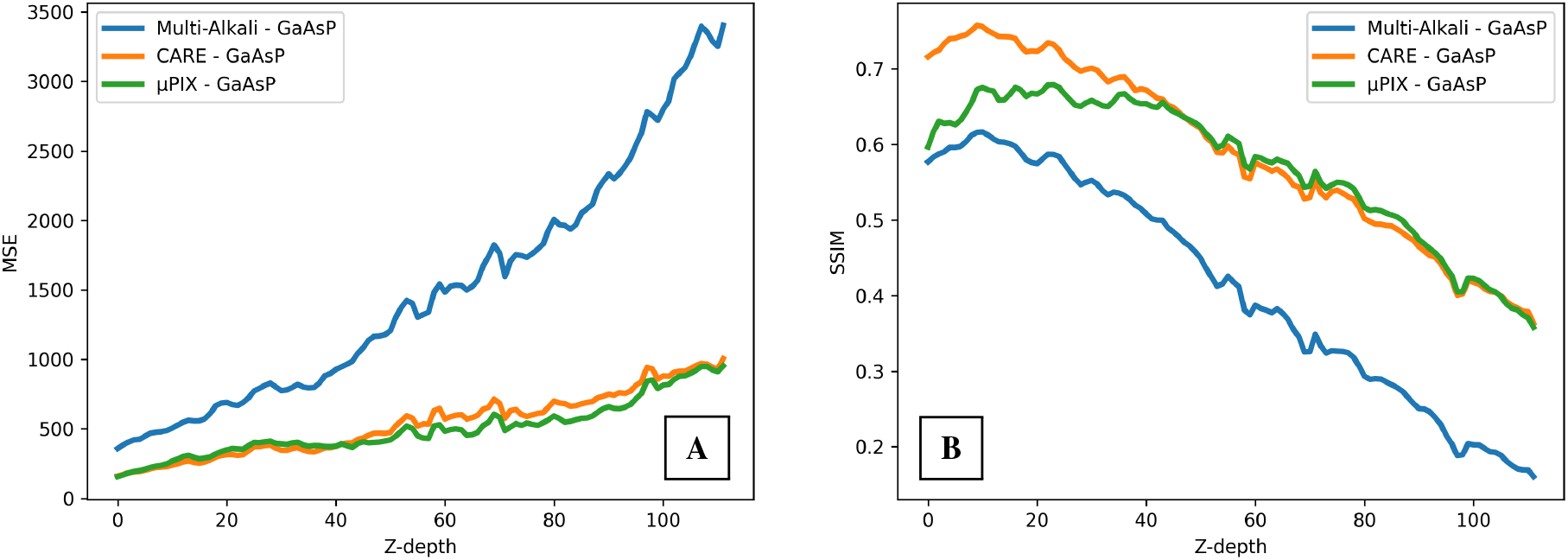
Evaluation of intensity and structural preservation along the Z-axis using the Microscope Rejuvenation Test stack. **A**: MSE between Multi-Alkali and GaAsP (blue), CARE and GaAsP (orange), *µPIX* and GaAsP (green) **B**: SSIM between Multi-Alkali and GaAsP (blue), CARE and GaAsP (orange), *µPIX* and GaAsP (green)

We next aimed to determine whether images generated by *µPIX* could be reliably used for quantitative imaging analysis. To do this, we evaluated whether *µPIX* maintains a consistent response across different levels of pixel intensity when compared to images acquired using GaAsP detectors. As shown in ***Figure 5A***, pixels ranging from 0 to nearly 60% of maximum intensity (representing approximately 95% of the total pixels in the test images) are restored with near-perfect linearity by *µPIX*. In contrast, ***Figure 5B*** shows that this consistency is not maintained by CARE. For pixel intensities greater than 60% and the maximum (representing around 5% of the image pixels), we observe moderate deviations from perfect restoration. However, this concerns only a small fraction of the pixels, most of which are saturated and therefore contain limited or non-informative data, making these deviations negligible for most practical purposes.

**Figure 5.**
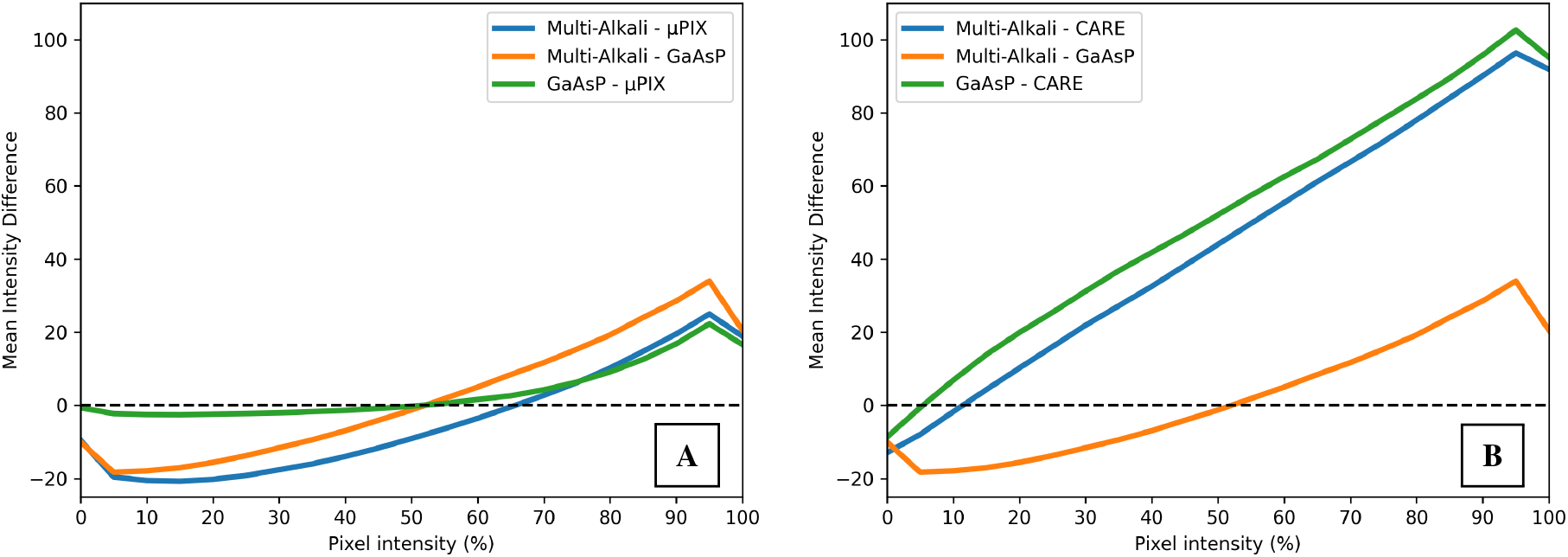
Evaluation of signal linearity preservation using the Microscope Rejuvenation Test Dataset. **A**: Mean intensity differences for pixel intensity ranging from 0 (no intensity) to 100 (maximum intensity) between Multi-Alkali and *µPIX* (blue), Multi-Alkali and GaAsP (orange), GaAsP and *µPIX*. The dashed line corespond to a perfect linearity. **B**: Mean intensity differences for pixel intensity ranging from 0 (no intensity) to 100 (maximum intensity) between Multi-Alkali and CARE (blue), Multi-Alkali and GaAsP (orange), GaAsP and CARE. The dashed line correspond to a perfect linearity.

This results suggests that using *µPIX* for detector rejuvenation opens the possibility for users to conduct quantitative imaging in much the same way as if they were using a GaAsP detector directly.

## DISCUSSION

A common concern among microscope users is the validity of AI generated images, as these are synthetic and may not be suitable for further analysis. The rise of generative AI has underscored the need to expand the family of perceptual metrics, focusing on human perception to validate such images. The two most widely used perceptual metrics are the Frechet Inception Distance (FID) and the Inception Score (IS) (25; 26). These metrics utilize a pre-trained Inception network on the ImageNet 10k dataset (27) and measure the Wasserstein distance and KL-divergence, respectively, between the embeddings of real and generated images. While these metrics are effective for well-structured images, they may be problematic in the context of image denoising. They tend to prioritize structural conservation over the preservation of overall distribution. Moreover, since these metrics rely on features extracted from a network pre-trained on a generalist images database, they may not be suitable for microscopic images. Microscopic images often contain subtle artifacts that may not align well with the features learned by the Inception model. To our knowledge, there is no perceptual metric specifically adapted to the nature of microscopic images that can effectively capture their unique feature space and accurately measure their perceptual quality. In line with previous attempts to assess image quality through human evaluation (28), we decided to adopt a similar approach. We designed an experiment involving 27 participants. These individuals were familiar with biological imaging but had no specific knowledge of the image categories or the methods used to generate them. Each participant completed seven rounds of evaluation, with each round consisting of four images: two generated by algorithms (*µPIX* and CARE) and two acquired through real detectors (Multi-Alkali and GaAsP), representing four distinct image categories. For each round, participants were asked to rate each image on a scale from 1 to 4, where 1 indicated the image was highly suitable for analysis, and 4 indicated it was completely unsuitable. The ratings for each image category were collected and their distributions are shown in ***Figure 6***. We conducted a statistical analysis between categories using the Wilcoxon non-parametric test. The evaluation results revealed that, as expected, a newer detector yields higher perceived image quality, as evidenced by a statistically significant distinction in human perception between images acquired with the older Multi-Alkali detector and those from the newer GaAsP detector (*p*-value = 7.87 *×* 10^−7^). We then compared the older Multi-Alkali images to both the CARE and *µPIX* generated images. Interestingly, in terms of human perception, the Multi-Alkali images and those enhanced by CARE were not judged to be significantly different, indicating that the enhancements applied by CARE do not improve the perceived usability of the images for analysis (*p*-value = 0.6151). In contrast, *µPIX* images were perceived as more usable for analysis when compared to Multi-Alkali(*p*-value = 6.63 *×* 10^−10^) and remarkably as more usable than those obtained from GaAsP detectors (*p*-value = 7.23 *×* 10^−3^). From a broader perspective, it appears that, despite its effectiveness in enhancing images, CARE (built on a convolutional autoencoder architecture) fails to convince users of its full applicability for image analysis. This is likely due to the known tendency of such networks to introduce blurring and smooth out structures (29; 30; 31). In contrast, generative approaches based on Pix2Pix networks aim to maintain the visual coherence of textures and structures while reducing noise. This leads to more realistic images that are better evaluated by human observers, and perceived similarly to real images captured by more advanced detectors.

**Figure 6.**
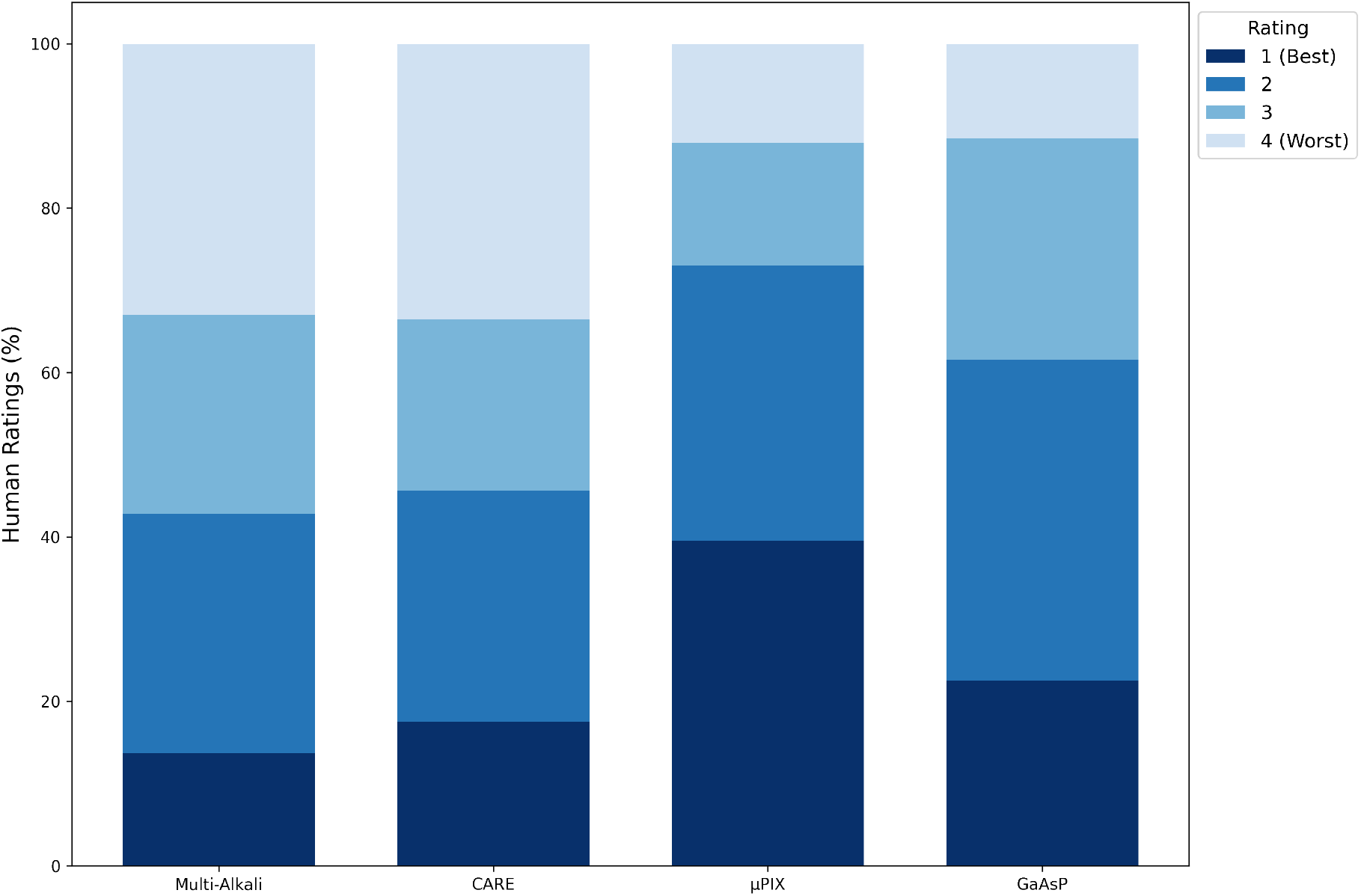
Evaluation of image quality by human observers. Stacked bar plot illustrating the distribution of human ratings for four image acquisition methods: Multi-Alkali, CARE, *µPIX*, and GaAsP. Each stacked bar represents the percentage of total ratings for each method, with the sum of ratings normalized to 100%. The ratings range from 1 (best: dark blue) to 4 (worst: light blue). The distribution reflects the overall quality perception across the different image types based on human evaluations.

Currently, the predominant paradigm for using Deep Learning in image treatment and analysis revolves around the use of U-Net networks, as seen in tools like CARE and CellPose3. The main advantages of these networks include their ease of training and deployment by non-specialist users, thanks to their training stability and the relatively short training time required to achieve good-quality results. This user-centered approach allows anyone to efficiently train their own model from scratch using their own dataset. However, we consider this approach unsustainable as it operates at the end of the imaging pipeline without addressing upstream factors such as hardware defects and specificities. Our *µPIX* workflow proposes an alternative paradigm, shifting from a user-centered to a hardware-centered perspective for model development and training. We envision that such a workflow should be developed over the long term by microscopy platforms to directly address hardware constraints and gradually build robust and flexible models for personalized microscopy. While it is well-known that training GANs has disadvantages compared to U-Nets such as longer training times (ranging from hours to days), training instability that can lead to mode collapse, and the challenge of determining an optimal stopping criterion; the results we present in terms of image denoising, segmentation, and hardware rejuvenation suggest that investing effort in developing models tailored to specific hardware by a deep learning specialist could be more beneficial. Theses models could then be shared by many users, as opposed to the current practice of many users developing rather identical models for the same hardware. In the era of *frugal AI*, we believe that our approach has the potential to save significant resources, both in terms of user time and global computational costs. Additionally, it is important to recognize that the ground truth datasets provided to *µPIX* are not of infinite quality and do not encompass every possible microscope artifact requiring correction. As the pix2pix architecture is based on a supervised learning approach, the system is inherently limited by the quality and variety of the images included in the training set. Consequently, *µPIX* cannot generate images of higher quality than those contained in the dataset, implying that it does not effectively correct hardware defects but rather learns to replicate them, producing images that closely resemble the provided ground truth.

Our approach opens up a wide range of possibilities for enhancing microscopy acquisition workflows and setups. Live imaging applications that require high prediction rates could particularly benefit from the use of *µPIX*. A major limitation of live imaging is phototoxicity, caused by the prolonged use of lasers. By utilizing a *µPIX* model pre-trained on a custom dataset with varying laser intensities, laser power can be reduced, potentially extending the lifespan of the samples being imaged and improving the image SNR. While the adversarial training process of Pix2Pix networks is time-consuming, the inference stage is relatively fast, as it only involves the network generator part, a lightweight UNET network. Benchmarking conducted on an Nvidia A6000 GPU with 48GB of VRAM revealed that *µPIX* can process 473 images of size 256×256 per second, 119 images of size 512×512, and 31 images of size 1024×1024. These results demonstrate that real-time image correction during live imaging sessions is greatly feasible, suggesting that such models can be seamlessly integrated into existing microscope setups. We have also begun exploring the application of *µPIX* for post-acquisition correction of temperature and oil refractive index mismatches.

We also explored whether an unsupervised approach could yield better results. The primary limitation of *µPIX* stems from the fact that Pix2Pix relies on a supervised learning architecture, which requires paired ground truth images registered at the same position for training. To address this, we investigated the use of *CycleGAN* architectures, which only require unpaired images from different conditions, making it easier to construct a training dataset. While the results were acceptable, the image quality was noticeably lower compared to what we achieved with Pix2Pix. We believe this limitation could be mitigated by developing a more robust and diverse dataset (***Supplementary Table 2***). Additionally, emerging models such as Diffusion networks and Transformer architectures, particularly Visual Transformers (ViT), represent a promising direction for future exploration, as they are rapidly becoming state-of-the-art for image generation and vision-related tasks.

In conclusion, our work introduces *µPIX* as an innovative solution to the pressing challenges of microscopy image denoising and restoration, effectively tackling issues arising from aging hardware and acquisition artifacts. Through rigorous comparisons with established denoising methods such as CARE and CellPose3, we have demonstrated that *µPIX* significantly enhances image quality, preserves structural integrity, and improves object segmentation accuracy. Our findings indicate that *µPIX* excels in rejuvenating images captured by older detectors, effectively bridging the gap to newer technologies while compensating for artifacts associated with light absorption along the Z-axis and maintaining a quasi-linear relationship in pixel intensity between the original and rejuvenated detector. By adopting a hardware-centered paradigm for model development, we emphasize the importance of creating specialized solutions tailored to specific microscopy setups, promoting a sustainable approach to image analysis that reduces computational burdens on users. Furthermore, our research underscores the transformative potential of *µPIX* in microscopy workflows, enabling researchers to confidently analyze synthetic images that closely mimic those obtained from advanced detectors. This work paves the way for future advancements and developments in generative AI for microscopy platforms, ultimately supporting the pursuit of more accurate and reliable imaging outcomes.

## MATERIAL AND METHODS

### Microscope setups

Fluorescence imaging was performed on a Zeiss LSM510 confocal scanning microscope (CLSM) equipped for two-photon imaging. The excitation laser is a Spectra Physics Mai Tai infrared tunable laser, used at 900 nm for the scope of these experiments. The scan head is mounted on a Zeiss Axiovert200M stand. We used a Plan-Apochromat 20x/0.75 Numerical Aperture objective. The room is well stabilized and controlled at a temperature of 21°C (*±* 1°C). The stand and sample environment was isolated in a black-painted incubation chamber, providing light isolation to prevent external signal pollution. The heating unit is not on, but the presence of the chamber still further helps with the temperature stability of the sample environment. Two beam paths were exploited for fluorescence detection. First, the more efficient non-descanned (NDD) beam path using two specially integrated Hamamatsu GaAsP detectors (new integration work in 2021 by ALPhANOV company). The detection range is set by filters from [500/550] nm for the *green* channel (the *red* channel was acquired but not used for the training). Second is the descanned configuration, using the internal Multi-Alkali PMT old detector of the scan head after the pinhole was fully opened. Filter sets of the internal beam path are chosen to fit a detection range as close as possible to the range of the NDD configuration of [500/550] nm. The system driving the acquisition software is Zeiss Efficient Navigation (ZEN) version 2009. To generate data without acquiring the same area, we use the tiling option with the overlap set to 0%.

### Two photons imaging

One-color two-photon imaging of immunostained samples was performed on an inverted Zeiss LSM510 confocal as described above. Multiposition imaging was used to automatically acquire image Z stacks on multiple gastruloids mounted on the same sample slide. The sampling parameters remain the same for all samples (pixel width 0.62 µm, voxel depth 1.2 µm), with 114 identical sections acquired sequentially on the GaAsP and Multi-Alkali detectors, reaching a depth in Z of 137 µm. The images were acquired with the full field-of-view (318 µm). The power of laser excitation and gains of detectors are optimized to exploit detector dynamics (8 bits) while avoiding any saturated pixels.

### Microscope Rejuvenation Dataset

#### Sample preparation

Gastruloids were generated using the protocol described previously in (24), from a H2B-GFP mouse embryonic stem cells line (a generous gift from Kat Hadjantonakis). Briefly, 200 cells were seeded and aggregated for 48 h in low-adherence 96-well plates (Costar ref: 7007) and subsequently pulsed with the Wnt agonist Chiron, which was washed out after 24 h, i.e. at 72 h of aggregate culture. We imaged 96 h old gastruloids, which exhibited polarized morphologies.

#### Dataset construction

As a result, we acquired 10 stacks of size 512×512 pixels with depth varying from 22 to 130 slices. Among those, we used 8 stacks to train/validate our model and two stacks to test the model. For the preparation of training and validation data, we tiled images into smaller patches of size 256×256 pixels with an overlap of 64 pixels to ensure comprehensive coverage. Employing a *reflect mode* for tile border padding helped mitigate any potential blank spaces resulting from the overlapping procedure. Furthermore, to standardize image intensity distributions, we applied percentile normalization between 1 and 99%, resulting in pixel values ranging from -1 to 1. During the training phase, we implemented data augmentation using the Albumentations (32) Python library. We decided to include transformations consistent with the biological objects used. We then used *shift scale rotate, elastic transformation, optical distorsion, randomrotate90* and *horizontal/vertical flip*.

### Loss and metrics

The training of the Pix2Pix model in this work is based on two primary loss functions: the adversarial loss and the L2 loss, which together guide the generator to produce realistic and accurate images. The adversarial loss is derived from a PatchGAN discriminator, which classifies each local patch of the image as real or fake, rather than making a single global decision for the entire image. The adversarial loss for the generator is defined as:

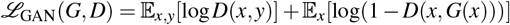

Here, *x* represents the input image, and *y* denotes the corresponding ground truth image. *D*(*x, y*) signifies the probability that the pair (*x, y*) is real, while *D*(*x, G*(*x*)) is the probability that the pair (*x, G*(*x*)) is generated by *G*. The generator seeks to minimize this loss, while the discriminator works to maximize it.

For both the discriminator and generator, we use *Binary Cross-Entropy (BCE)* loss to quantify their performance. The discriminator’s loss is computed as:

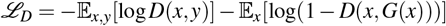

where the discriminator aims to correctly classify real patches from the ground truth *y* and generated patches from *G*(*x*).

In this work, instead of using an L1 loss, the generator is trained using an *L2 loss*, or mean squared error (MSE), to penalize pixel-wise differences between the generated image and the ground truth image. The L2 loss is expressed as:

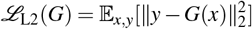

The L2 loss encourages the generator to produce outputs that closely match the target image at the pixel level by minimizing the squared differences between corresponding pixel values. This loss emphasizes structural accuracy and helps prevent large deviations in pixel intensities.

In our setup, instead of using a single balancing hyperparameter *λ*, we employed two separate weight parameters, *w*_1_ and *w*_2_, to control the contribution of the adversarial loss and the reconstruction loss, respectively. The overall objective function for the generator is therefore formulated as:

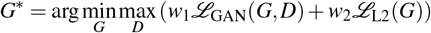

Here, *w*_1_ controls the influence of the adversarial loss, ensuring that the generated images are realistic enough to fool the discriminator, while *w*_2_ controls the contribution of the L2 reconstruction loss, ensuring that the generated images closely match the target images at a pixel level. The combination of these two losses allows the generator to produce both perceptually realistic and structurally accurate images.

### Metrics

For image denoising, we employ the *Mean Squared Error (MSE)* and *Structural Similarity Index (SSIM)*. For segmentation quality assessment, we utilize *Intersection over Union (IoU), Precision, Recall*, and *F1-Score*.

#### Mean Squared Error (MSE)

The MSE quantifies the average squared difference between the actual and predicted values. It is calculated using the following formula:

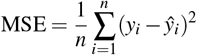

In this equation, *y*_*i*_ represents the true value, *ŷ*_*i*_ denotes the predicted value, and *n* is the number of data points. MSE measures the average of the squares of the errors, indicating how closely the predicted values match the true values.

#### Structural Similarity Index (SSIM)

The SSIM evaluates the perceptual similarity between two images, considering factors such as luminance, contrast, and structure. The SSIM is computed as follows:

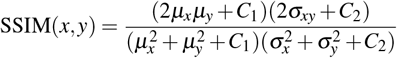

Here, *µ*_*x*_ and *µ*_*y*_ are the average pixel values of images *x* and *y*, 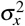 and 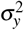 are the variances of images *x* and *y*, and *σ*_*xy*_ is the covariance between *x* and *y*. The constants *C*_1_ and *C*_2_ are used to stabilize the division with weak denominators.

#### Intersection over Union (IoU)

Intersection over Union (IoU) measures the overlap between the predicted segmentation mask and the ground truth mask. It is defined by the following formula:

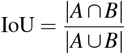

In this context, *A* represents the set of pixels in the predicted segmentation mask, while *B* denotes the set of pixels in the ground truth mask. The term |*A*∩*B*| refers to the number of pixels in the intersection of the predicted and ground truth masks, and |*A*∪*B*| refers to the number of pixels in the union of these masks.

#### Precision

Precision measures the accuracy of positive predictions. It is calculated using the formula:

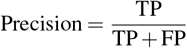

In this equation, TP stands for the number of true positives, while FP denotes the number of false positives.

#### Recall

Recall quantifies the model’s ability to identify all relevant positive instances. It is given by:

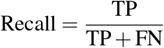

Here, TP represents the number of true positives, and FN indicates the number of false negatives.

#### F1-Score

The F1-score is the harmonic mean of Precision and Recall, providing a single metric that balances both aspects. It is defined as:

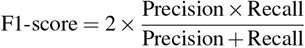

### *µPIX* Architecture, Training and Inference

We employed an architecture based on the Pix2Pix, consisting of a generator that leverages the TensorFlow Segmentation Model library (33) using a classical U-Net architecture with a non-pretrained EfficientNet-B0 backbone. The discriminator was based on a traditional convolutional PatchGAN network, which classifies image patches of size 16×16 using a binary classifier with a sigmoid activation function in the output layer. Input images were tiled from the original dataset into 256×256 patches, and the generated output images matched this size. Pixel intensities in the output were remapped to the range [−1, 1] using a hyperbolic tangent activation function in the generator’s final layer.

During training, the generator aimed to minimize the dissimilarity between the generated images and their corresponding ground truth counterparts, while the discriminator sought to differentiate between real and generated images. This adversarial training process facilitated a continuous cycle, wherein the generator progressively refined its output to appear more realistic, and the discriminator became increasingly adept at distinguishing real from generated images. This iterative optimization enabled the generator to produce high-fidelity image translations closely resembling the ground truth, while the discriminator enhanced its classification accuracy.

To optimize the model, we utilized binary cross-entropy (BCE) loss for the discriminator and MSE for the generator. The overall GAN loss was implemented as a weighted combination of these losses, with the BCE weight (w1) set to 10 and the MSE weight (w2) set to 100. Both components were trained with a constant learning rate of 1 *×* 10^−4^ using the Adam optimizer in TensorFlow. Our models were trained on an NVIDIA A6000 GPU with 48 GB of memory, utilizing a batch size of 128 to efficiently process large datasets.

Since determining optimal stopping criteria for generative adversarial networks is challenging, we implemented a custom early stopping method. At the end of each epoch, the ratio between the MSE and the SSIM was evaluated on the validation set. If this ratio reached a new minimum, the model was saved, and training continued for a set number of epochs defined by the patience hyperparameter (set to 20 by default). If no improvement was observed within the patience period, training was halted. During inference, only the generator component was used, and the pixel range of the generated images was remapped to [0, 255] for consistency with the original input image format.

### *µPIX* Software

#### Installation

*µPIX* is a Python package composed of three primary scripts: `training.py`, which facilitates training from scratch or retraining/fine-tuning a model; `predict.py`, which performs inference on new images using a pretrained model; and `new experiment.py`, which initializes and organizes µPIX experiments. The software can be installed by either cloning the GitHub repository or by using the pre-built Docker image (*See Code, Data and model availability* section)

#### Workflow

The µPIX workflow revolves around the concept of “experiments.” Each experiment is initialized using the `new experiment.py` script, which defines the experiment’s name, location, and the directories containing the images used for training and testing. This script generates an `experiment directory` that includes a `hyperparameters.json` file, where user-defined and default *µPIX* hyperparameters (e.g., `batch size`, `learning rates`, etc.) are stored for subsequent use.

~~~
python new_experiment.py --name my_experiment
                         --path ./experiments/my_experiment
                         --noisy-image-path ./data/noisy
                         --clean-image-path ./data/clean
                         --test-image-path ./data/test
~~~

Following the experiment setup, model training is initiated using the `training.py` script. This script reads the `hyperparameters.json` file and creates a `Results` directory within the experiment folder.

The `Results` directory contains the trained network, validation images generated during training, and a log file (`log.txt`) that tracks the training progress and validation loss.

~~~
python training.py --experiment-path ./experiments/my_experiment
~~~

Additionally, users can retrain or fine-tune a previously trained model by using the `--retrain` option, allowing for further optimization or adjustment of the model based on new data or parameters:

~~~
python training.py --experiment-path ./experiments/my_experiment --retrain
~~~

Once the training is completed, the `predict.py` script is employed to perform inference on new data store into the test folder defined in the experiment. The predictions are stored in a `Predictions` directory, which is automatically generated within the experiment folder.

~~~
python predict.py --experiment-path ./experiments/my_experiment
~~~

## Code, Data and model availability

- Code & Data: https://giltab.lis-lab.fr/sicomp/muPIX.

## ACKNOWLEDGMENTS

We acknowledge the France-Bioimaging Infrastructure (ANR-10-INBS-04). We would like to thank the GDR Imabio consortium (CNRS) for funding the start of this project with an exploratory master’s fellowship and the IBDM management and CNRS for funding Gabriel Bon. Special thanks to Sham Tlili for providing us with gastruloids samples and stimulating discussions. We thank all IBDM members who join us to contribute to image-blind evaluation.

## REFERENCES

[1] H. Lanteri, C. Aime, H. Beaumont, and P. Gaucherel, “Blind deconvolution using the Richardson-Lucy algorithm,” in Optics in Atmospheric Propagation and Random Phenomena, A. Kohnle and A. D. Devir, Eds., vol. 2312, International Society for Optics and Photonics. SPIE, 1994, pp. 182 – 192. [Online]. Available: 10.1117/12.197374

[2] L. Chen, J. Zhang, Z. Li, Y. Wei, F. Fang, J. Ren, and J. Pan, “Deep richardson-lucy deconvolution for low-light image deblurring,” 2023. [Online]. Available: https://arxiv.org/abs/2308.05543

[3] J. Chen, J. Benesty, Y. Huang, and S. Doclo, “New insights into the noise reduction wiener filter,” IEEE Transactions on Audio, Speech, and Language Processing, vol. 14, no. 4, pp. 1218–1234, 2006.

[4] C. Bled and F. Pitiè, “Pushing the limits of the wiener filter in image denoising,” 2023. [Online]. Available: https://arxiv.org/abs/2303.16640

[5] A. H. Klemm, A. W. Thomae, K. Wachal, and S. Dietzel, “Tracking Microscope Performance: A Workflow to Compare Point Spread Function Evaluations Over Time,” Microscopy and Microanalysis, vol. 25, no. 3, pp. 699–704, 06 2019. [Online]. Available: 10.1017/S1431927619000060

[6] N. Gritti, R. M. Power, A. Graves, and J. Huisken, “Image restoration of degraded time-lapse microscopy data mediated by near-infrared imaging,” Nature Methods, vol. 21, no. 2, pp. 311–321, Feb 2024. [Online]. Available: 10.1038/s41592-023-02127-z

[7] M. Weigert, U. Schmidt, T. Boothe, A. Müller, A. Dibrov, A. Jain, B. Wilhelm, D. Schmidt, C. Broaddus, S. Culley, M. Rocha-Martins, F. Segovia-Miranda, C. Norden, R. Henriques, M. Zerial, M. Solimena, J. Rink, P. Tomancak, L. Royer, F. Jug, and E. W. Myers, “Content-aware image restoration: pushing the limits of fluorescence microscopy,” Nature Methods, vol. 15, no. 12, pp. 1090–1097, Dec 2018. [Online]. Available: 10.1038/s41592-018-0216-7

[8] A. Krull, T.-O. Buchholz, and F. Jug, “Noise2void - learning denoising from single noisy images,” 2019. [Online]. Available: https://arxiv.org/abs/1811.10980

[9] J. Lehtinen, J. Munkberg, J. Hasselgren, S. Laine, T. Karras, M. Aittala, and T. Aila, “Noise2noise: Learning image restoration without clean data,” 2018. [Online]. Available: https://arxiv.org/abs/1803.04189

[10] C. Stringer, T. Wang, M. Michaelos, and M. Pachitariu, “Cellpose: a generalist algorithm for cellular segmentation,” Nature Methods, vol. 18, no. 1, pp. 100–106, Jan 2021. [Online]. Available: 10.1038/s41592-020-01018-x

[11] M. Pachitariu and C. Stringer, “Cellpose 2.0: how to train your own model,” Nature Methods, vol. 19, no. 12, pp. 1634–1641, Dec 2022. [Online]. Available: 10.1038/s41592-022-01663-4

[12] C. Stringer and M. Pachitariu, “Cellpose3: one-click image restoration for improved cellular segmentation,” bioRxiv, 2024. [Online]. Available: https://www.biorxiv.org/content/early/2024/02/12/2024.02.10.579780

[13] R. Zhou, M. E. Helou, D. Sage, T. Laroche, A. Seitz, and S. Süsstrunk, “W2s: Microscopy data with joint denoising and super-resolution for widefield to sim mapping,” 2020. [Online]. Available: https://arxiv.org/abs/2003.05961

[14] M. Chatton, “Microscopy image restoration using deep learning on w2s,” 2020. [Online]. Available: https://arxiv.org/abs/2004.10884

[15] Y. Wu, Y. Rivenson, H. Wang, Y. Luo, E. Ben-David, L. A. Bentolila, C. Pritz, and A. Ozcan, “Three-dimensional virtual refocusing of fluorescence microscopy images using deep learning,” Nature Methods, vol. 16, no. 12, pp. 1323–1331, Dec 2019. [Online]. Available: 10.1038/s41592-019-0622-5

[16] O. Ronneberger, P. Fischer, and T. Brox, “U-net: Convolutional networks for biomedical image segmentation,” in Medical Image Computing and Computer-Assisted Intervention – MICCAI 2015, N. Navab, J. Hornegger, W. M. Wells, and A. F. Frangi, Eds. Cham: Springer International Publishing, 2015, pp. 234–241.

[17] M. Tan and Q. V. Le, “Efficientnet: Rethinking model scaling for convolutional neural networks,” 2020. [Online]. Available: https://arxiv.org/abs/1905.11946

[18] C. Li and M. Wand, “Precomputed real-time texture synthesis with markovian generative adversarial networks,” 2016. [Online]. Available: https://arxiv.org/abs/1604.04382

[19] I. J. Goodfellow, J. Pouget-Abadie, M. Mirza, B. Xu, D. Warde-Farley, S. Ozair, A. Courville, and Y. Bengio, “Generative adversarial networks,” 2014. [Online]. Available: https://arxiv.org/abs/1406.2661

[20] V. Ljosa, K. L. Sokolnicki, and A. E. Carpenter, “Annotated high-throughput microscopy image sets for validation,” Nature Methods, vol. 9, no. 7, pp. 637–637, Jul 2012. [Online]. Available: 10.1038/nmeth.2083

[21] U. Schmidt, M. Weigert, C. Broaddus, and G. Myers, Cell Detection with Star-Convex Polygons. Springer International Publishing, 2018, p. 265–273. [Online]. Available: 10.1007/978-3-030-00934-230

[22] Hamamatsu, “Photomultiplier tube,” 2024. [Online]. Available: https://www.hamamatsu.com/us/en/product/optical-sensors/pmt/pmttube-alone/side-on-type/R6357.html

[23] Hamamatsu1, “H7422p-40,” 2024. [Online]. Available: https://www.alldatasheet.com/datasheet-pdf/pdf/62585/HAMAMATSU/H7422P-40.html

[24] A. Hashmi, S. Tlili, P. Perrin, M. Lowndes, H. Peradziryi, J. M. Brickman, A. Martínez Arias, and P.-F. Lenne, “Cell-state transitions and collective cell movement generate an endoderm-like region in gastruloids,” eLife, vol. 11, apr 2022. [Online]. Available: https://elifesciences.org/articles/59371

[25] M. Heusel, H. Ramsauer, T. Unterthiner, B. Nessler, and S. Hochreiter, “Gans trained by a two time-scale update rule converge to a local nash equilibrium,” 2018. [Online]. Available: https://arxiv.org/abs/1706.08500

[26] S. Barratt and R. Sharma, “A note on the inception score,” 2018. [Online]. Available: https://arxiv.org/abs/1801.01973

[27] J. Deng, W. Dong, R. Socher, L.-J. Li, K. Li, and L. Fei-Fei, “Imagenet: A large-scale hierarchical image database,” in 2009 IEEE Conference on Computer Vision and Pattern Recognition, 2009, pp. 248–255.

[28] K. Combs, T. J. Bihl, A. Gadre, and I. Christopherson, “A human-factors approach for evaluating ai-generated images,” in Proceedings of the 2024 Computers and People Research Conference, ser. SIGMIS-CPR ‘24. New York, NY, USA: Association for Computing Machinery, 2024. [Online]. Available: 10.1145/3632634.3655849

[29] H. Hudson and T. C. Lee, “Maximum likelihood restoration and choice of smoothing parameter in deconvolution of image data subject to poisson noise,” Computational Statistics and Data Analysis, vol. 26, no. 4, pp. 393–410, 1998. [Online]. Available: https://www.sciencedirect.com/science/article/pii/S0167947397000418

[30] N. Ichimura, “Spatial frequency loss for learning convolutional autoencoders,” ArXiv, vol. abs/1806.02336, 2018.

[31] S. Hong and S.-K. Song, “Kick: Shift-n-overlap cascades of transposed convolutional layer for better autoencoding reconstruction on remote sensing imagery,” IEEE Access, vol. 8, pp. 107 244–107 259, 2020.

[32] K. E. I. V. K. A. Buslaev A, Parinov A, “Albumentations: fast and flexible image augmentations,” ArXiv e-prints, 2018.

[33] P. Iakubovskii, “Segmentation models,” https://github.com/qubvel/segmentationmodels, 2019.

