## Supplemental Data for "*µPIX*: Leveraging Generative AI for Enhanced, Personalized and Sustainable Microscopy"

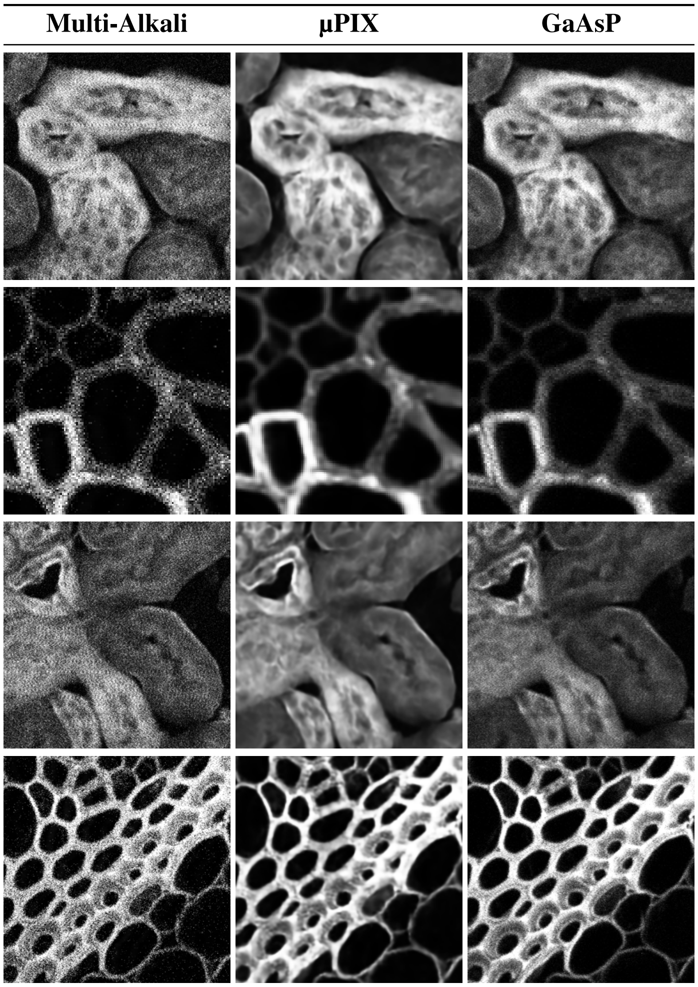


**Table 1.** ***µPIX* denoising results on a metrology dataset.**
Some results obtained with a *µPIX* model trained on a custom dataset consisting of various biological objects and textures : convalaria and mouse kidney.

| Multi-Alkali | CycleGAN | GaAsP |
|---|---|---|

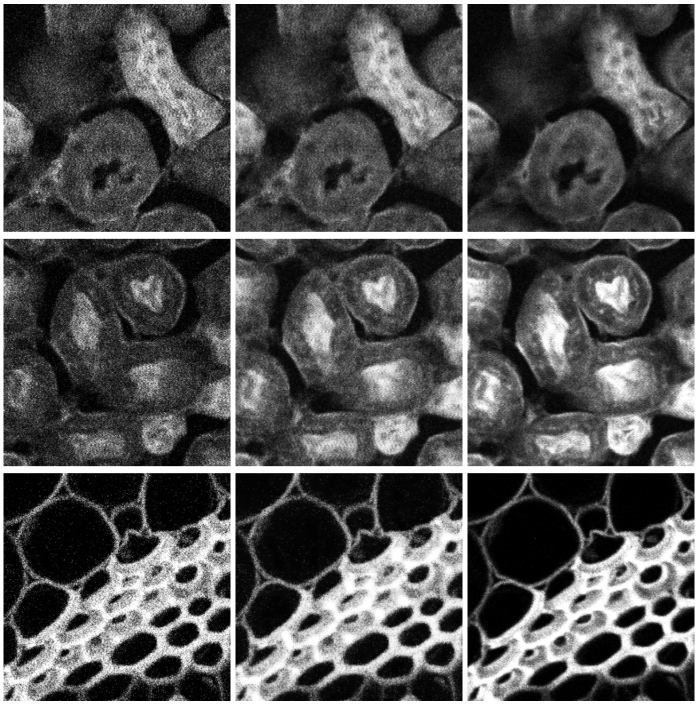

**Table 2.** ***CycleGAN* denoising results on a metrology dataset .**
Some results obtained with a *CycleGAN* model trained on a custom dataset consisting of various biological objects and textures : convalaria and mouse kidney.
